# Species-dependent effects of seawater acidification on alkaline phosphatase activity in dinoflagellates

**DOI:** 10.1101/2022.12.10.519933

**Authors:** Chentao Guo, Ling Li, Senjie Lin, Xin Lin

## Abstract

Global climate change is widely shown to cause ocean acidification (OA) and projected to intensify nutrient limitation. Phosphorus (P) is an essential nutrient for phytoplankton to grow. While dissolved inorganic phosphate (DIP) is the preferred form of P, phytoplankton have evolved alkaline phosphatase (AP) to utilize dissolved organic phosphorus (DOP) when DIP is deficient. Although AP is known to require pH>7, how OA may affect AP activity and hence the capacity of phytoplankton to utilize DOP is poorly understood. Here, we examine the effects of pH conditions (5.5 to 11) on AP activity from six species of dinoflagellates, an important group of marine phytoplankton. We observed a general pattern that AP activity declined sharply at pH 5.5, peaked between pH 7 and 8, and dropped at pH>8. However, our data revealed remarkable interspecific variations in optimal pH and niche breadth of pH. Among the species examined, *Fugacium kawagutii* and *Prorocentrum cordatum* had an optimal pH at 8, *Alexandrium pacificum, Amphidinium carterae, Effrenium voratum*, and *Karenia mikimotoi* showed an optimal pH of 7. However, whereas *A. pacificum, F. kawagutii* and *K. mikimotoi* had a broad pH range for AP (7-11), *A. carterae, E. voratum*, and *P. cordatum* exhibited a narrow pH range. The response of AP activity of *A. carterae* to pH changes was verified using purified AP heterologously expressed in *Escherichia coli*. These findings suggest that OA will likely differentially impact the capacity of different phytoplankton species to utilize DOP in the projected acidified and nutrient-limited future ocean.

## 1. Introduction

Due to fossil fuel consumption, atmospheric CO_2_ has rapidly increased from the pre-industrial 280 ppm to the current ∼410 ppm and is projected to reach 700-1000 ppm at the end of the century (Alley et al. 2007). As approximately 30% of this anthropogenically emitted CO_2_ is taken up by the oceans (Feely et al. 2004, Sabine et al. 2004), one major consequence of rising CO_2_ is ocean acidification (OA), which manifests as the decline of seawater pH. Over the last 250 years, the average pH of the ocean has dropped from about 8.2 to 8.1, which is approximately equivalent to a 30% increase in H^+^ concentration (Doney et al. 2009). Existing models predict that during the 21^st^ century, surface ocean pH may decline by additional 0.3–0.4 units (Orr et al. 2005, Caldeira and Wickett 2005). Phytoplankton will be the first responders to OA, because their photosynthesis interacts intimately with changing CO_2_ and pH conditions (Rose et al. 2009, Hutchins et al. 2019).

Effect of OA is not even among phytoplankton species. Studies have shown that OA promotes growth of some phytoplankton species and inhibits growth of others and has no detectable effect on yet other species (Dutkiewicz et al. 2015, Gao et al. 2019). This variability is due to the distinct genetic and physiology of individual species of phytoplankton. It is believed that OA can benefit species that usually rely on energetically expensive carbon concentrating mechanisms (CCM) to obtain adequate CO_2_ for photosynthesis, because Rubisco is less than one-half saturated under current CO_2_ levels (Giordano et al. 2005). However, OA also stresses phytoplankton, as there is an energetic cost to maintain cellular pH homeostasis. The net effect therefore is dependent on the offset between the beneficial and the stressful effects of OA. Hinga reviewed historical data suggesting that changes in seawater pH could affect differentially affect different phytoplankton species in terms of growth rates and abundance, and hence changes in phytoplankton communities (Hinga 2002).

OA does not act alone but can interact with nutrient dynamics to impact phytoplankton. Another consequence of rising atmospheric CO_2_ is global warming, which intensifies water column stratification and impedes upwelling of nutrient-rich deep waters. Therefore, OA may be compounded by nutrient deficiency in the future ocean. Besides, excessive nitrogen loading in coastal waters can cause stoichiometric deficiency in phosphorus (P). P is an essential nutrient for the growth of phytoplankton. In the ocean, P occurs as dissolved inorganic phosphate (DIP) and dissolved organic phosphate (DOP). DIP is the preferred form for phytoplankton and usually is depleted in the surface ocean (Benitez-Nelson 2000). In contrast, DOP is relatively abundant in the eutrophic zone. When facing the DIP deficiency, phytoplankton can utilize DOP, mainly due to alkaline phosphatase (AP) that phytoplankton (as well as bacteria) have evolved (Dyhrman and Ruttenberg 2006, Lin et al. 2016).

As AP is believed to function optimally at basic pH (>7), OA can potentially influence AP activity and hence phytoplankton’s capacity to utilize DOP. However, the potential effect is poorly understood and severely underexplored. There are diverse types of AP in phytoplankton, with substantial sequence divergence between species of phytoplankton. New types of AP have continually been discovered (e.g. Dyhrman and Palenik 1997, Xu et al. 2006, Lin et al. 2011, Lin et al. 2012a, Lin et al. 2012b, Sun et al. 2012, Lin et al. 2013). Conceivably, these APs may respond differently to OA, but experimental evidence is limited.

In this study, we investigated the effect of pH on APs in six dinoflagellate species (*Amphidinium carterae, Alexandrium pacificum, Karenia mikimotoi, Prorocentrum cordatum* [formerly *P. minimum*], *Fugacium kawagutii and Efferenium voratum*). To better demonstrate the pH dependency of phytoplankton APs, we overexpressed the AP *A. carterae* (ACAAP) in *Escherichia coli* and conducted activity assay on purified recombinant ACAAP (rACAAP).

## 2. Materials and Methods

### 2.1 Algal Strains, Culture Conditions and Sample Collection

Six dinoflagellate strains were used in this study. *A. carterae and F. kawagutii* were provided by the Provasoli-Guillard National Center for Marine Algae and Microbiota (NCMA); E. *voratum*, A. *pacificum* and *P. cordatum (*formerly *P. minimum*) were provided by Center for Collections of Marine Bacteria and Phytoplankton, Xiamen University (CCMBP); and *K. mikimotoi* was provided by Jinan University. These strains were maintained in sterilized oceanic seawater (filtered through 0.22 μm pore size filters, salinity 30) amended with the full nutrient regime of the L1 medium. Temperature was set at 20°C except for *F. kawagutii*, which was grown at 25 °C. Illumination was provided under a light dark cycle L:D = 12:12 with a photon flux of 100 μE m^-2^ s^-1^.

In experiments, DIP-depleted and replete cultures were prepared in triplicates. The DIP-depleted cultures were grown under the same nutrient regime as DIP-replete cultures, except for a reduced phosphate concentration (2 μM instead of 36 μM). Temperature and light conditions were the same described above. The cell density in the cultures was determined daily by cell count carried out using a Sedgewick–Rafter counting chamber (Phycotech, St. Joseph, MI, USA). The DIP concentration in the culture media was measured using the Molybdenum Blue method (Parsons et al. 1984).

### 2.2 Heterologous Expression of Recombinant ACAAP

To determine direct effects of pH on AP enzyme, AP of *A. carterae* was expressed in *Escherichia coli* and purified. Based on the AP-encoding gene sequence (*Acaap*, GenBank: HQ259111.2) identified in *A. carterae* (Lin et al.2011), a recombinant ACAAP (rACAAP) comprising 684 amino acid residues (full ORF region with the exclusion of a signal peptide coding region (nucleotide site 1-60)) was used. Gene fragments were amplified using specific primers PF2 (5’-AAGGGACGTAGGCTTGCTAG-3’) and PR2 (5’-CGCACGCACGGTCAAGAAG - 3’). PCR was carried out in 25 µL composed of 1X PCR buffer, 2µL (10 mmol/L) each nucleotide, 1µL (5 μmol/L) each primer, 1µL template DNA, and 0.75 unit Taq under the program consisting of the denaturing step at 94°C for 3min, followed by 35 cycles of 94°C for 30s, 56°C for 45s, and 72°C for 1 min, and ended with a final extension at 72°C for 10 min. The PCR product was gel-purified and cloned into the expression vector pEasy-E1 (TransGen Biotech, Beijing, China) and transformed into *E. coli* BL21 (DE3) (TransGen Biotech, Beijing, China). A single colony containing the gene fragment was grown in a 3 mL LB (Luria-Bertani) liquid medium containing 100 μg/mL ampicillin on a shaker rotating at 200 rpm at 37°C for 16 h; each colony was then transferred to 1 L fresh LB with 0.5 mM IPTG (isopropyl-β-D-thiogalactopyranoside) to induce the expression of the inserted *Acaap* gene fragment, grown for10 h at 28°C and shaken at 200 rpm.

*E. coli* cells were harvested by centrifugation at 5000 x g at 4°C for 10 min, resuspended in a 1mL Tris-HCl buffer (50 mM Tris-HCl, pH 8.0), and then homogenized by ultrasonic treatment. After centrifugation at 15,000 x g, 4°C for 10min, the supernatant containing the heterologously expressed peptide was purified using an Ni-NTA spin kit column (TransGen Biotech, Beijing, China) following the manufacturer’s protocol. The resultant overexpressed peptide was loaded into a Superdex 75 gel filtration column on an AKTA prime liquid chromatography system (GE Healthcare Bio-Sciences, Uppsala, Sweden) and eluted using a phosphate-buffered saline buffer (PBS buffer, 50 mM NaH_2_PO_4_, 150mM NaCl, pH 8.0) as the mobile phase. All collected fractions were examined by sodium dodecyl sulfate polyacrylamide gel electrophoresis (SDS-PAGE), and fractions containing the target peptide were concentrated using Amicon Ultra centrifugal filter devices (Merck Millipore Ltd., Carrigtwohill, IRL). Purified rACAAP was subjected to further enzymatic characterization.

### 2.3 pH Dependency Analysis of AP activity

The pH dependency analysis of AP was investigated in six dinoflagellate species and the purified rACAAP. The dinoflagellate species were grown under the DIP-depleted condition as described above, and ∼10^6^ cells were collected at the stationary phase using centrifugation at 5000 x g at the same temperature as cultures for 10 min. Cell pellets were resuspended in PBS (pH 7.4) and homogenized using 0.5 mm ceramic beads beating with the setting of 6 m s^−1^ for 1min by MP FastPrep-24 (MP Biomedicals, CA, USA). After centrifugation at 13000 x g for 2 min at 20°C, total protein in the supernatant was removed to a clean tube and protein concentration was measured using BCA (Tiangen Research, Beijing, China). Six μg protein from each sample was used for AP activity assays, which contained 75μL AP buffer (50mM Tris-Cl, 20 mM NaCl) with a pH gradient (5.5-11.0), 5 μL Ca^2+^ (100 mmol/L), 5 μL Mg^2+^ (100 mmol/L), 5 μL 20 mM pNPP. The reactions were incubated at 25°C in darkness for 2 hours in a 96-well plate. Triplicate wells for each sample were measured.

## 3. Results and Discussion

The effect of pH assay was examined on crude protein extracted from the six dinoflagellate species included in this study, which were grown under the P-depleted condition to induce AP expression. As shown in Fig. 1, there was a strong response in AP activity to pH changes in general, and the mode of response varied with species. At low pH (5.5), AP activity was the lowest in all six species. Half of the species exhibited the highest AP activity at pH 7 (*A. carterae, E. voratum*, and *K. mikimotoi*; Fig. 1b, c, e), whereas two species showed the highest level of AP activity at pH 8 (*F. kawagutii* and *P. cordatum*; Fig. 1d, f). Three species displayed a broad range of suitable pH (7-10 for *A. pacificum*; 7-11 for *K. mikimotoi*; 8-11 for *F. kawagutii*; Fig. 1a, e, d) whereas the other three showed declined AP activity at pH>7 or 8 (Fig. 1b, c, f).

**Fig. 1.**
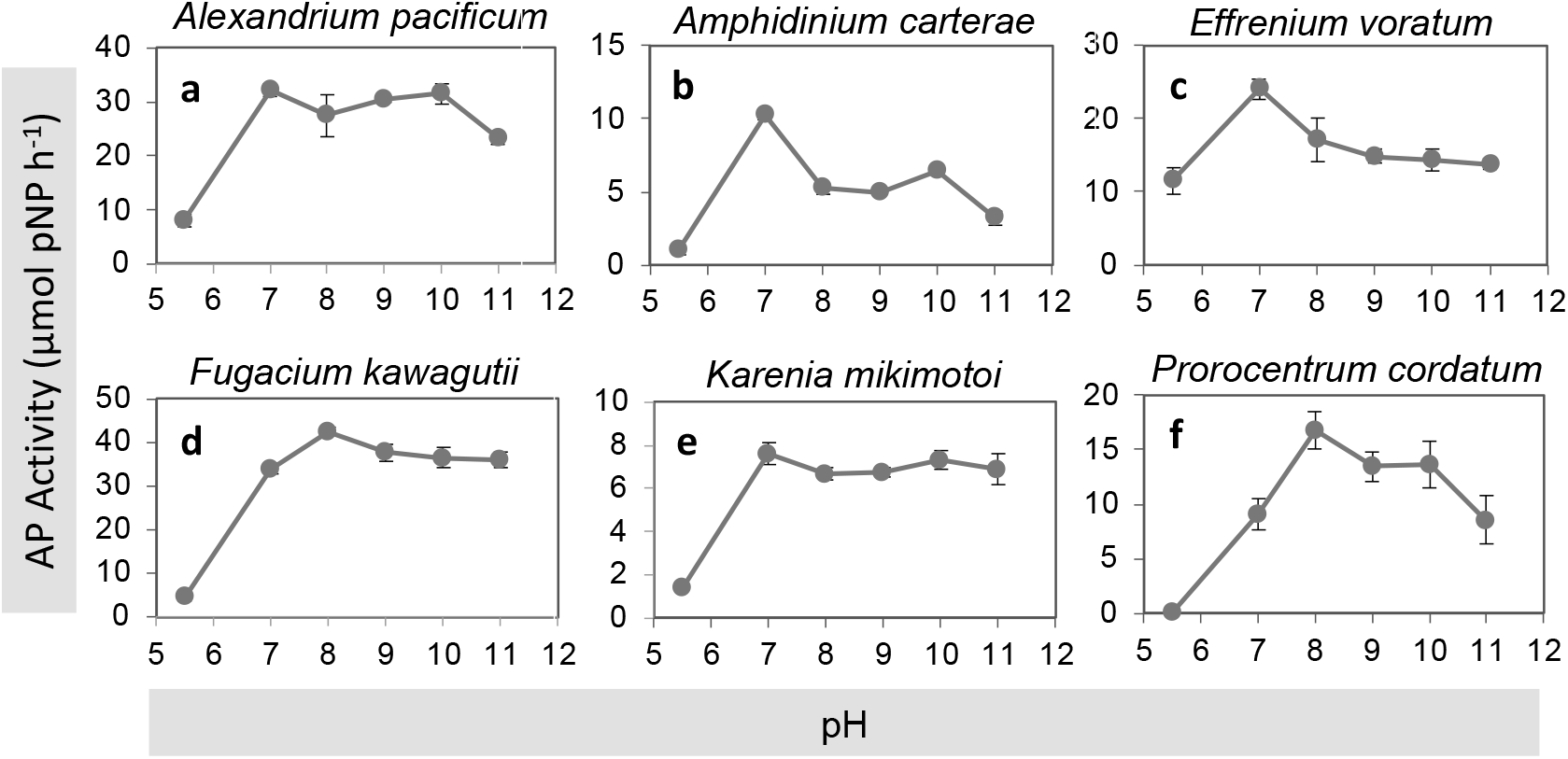
The effect of pH on alkaline activity in different species of dinoflagellates.

To verify that the responses of AP activity to pH variation was directly from the AP enzyme, we expressed AP in *A. carterae* (ACAAP) in *E. coli* and purified rACAAP (Fig. 2). The same AP activity assay as for crude proteins was conducted on the purified rACAAP. The result (Fig. 3) was consistent with the observation of crude protein extracts of *A. carterae* (Fig. 1b), in both of which the optimum pH was 7.

**Fig. 2.**
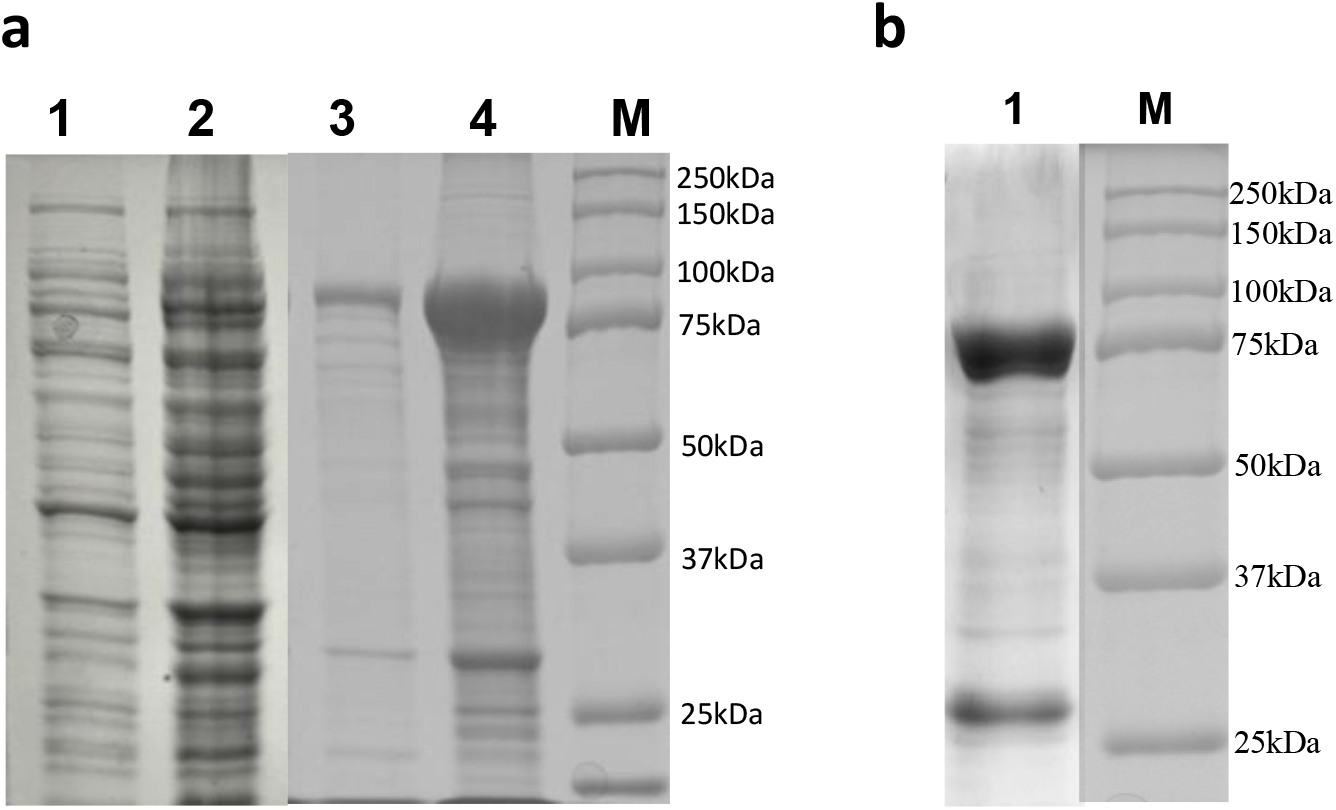
SDS-PAGE analysis of rACAAP expressed in *E. coli*. **a**. Analysis of crude protein extract. Lanes: M, Bio-Rad Protein Marker; 1, Pre-induced recombinant protein; 2, After-induced recombinant protein; 3, Cell ultrasound sediment; 4, Cell ultrasound supernatant. The thick band between 75 and 100 kDa is rACAAP. **b**. Analysis of purified rACAAP. Lanes: M, Bio-Rad Protein Marker; 1, Purified rACAAP (band between 75 and 100 kDa).

**Fig. 3.**
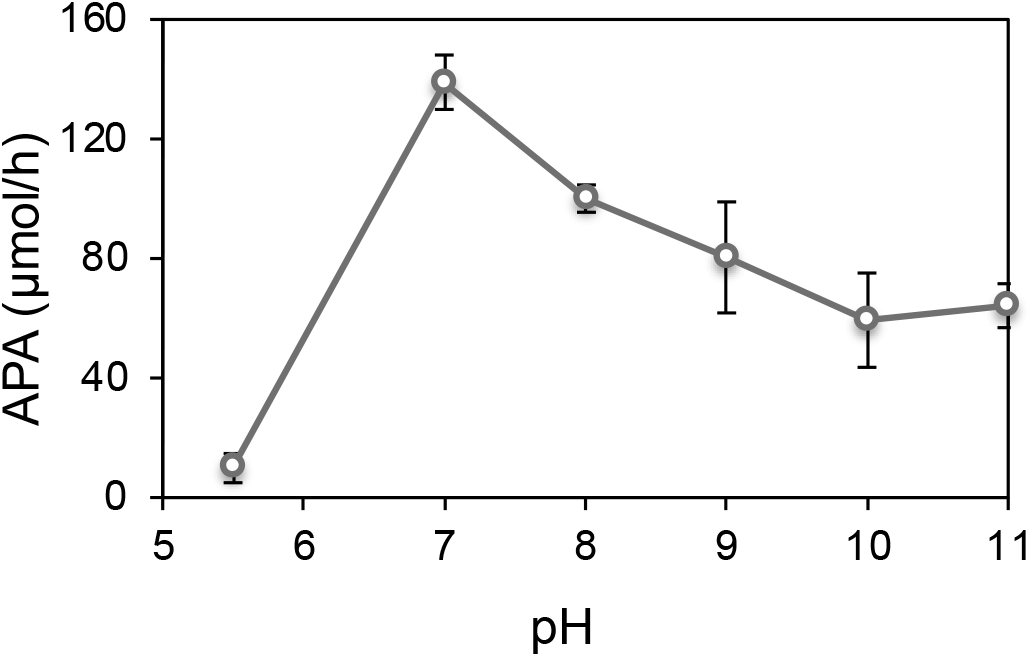
The effect of pH on AP activity of purified rACAAP.

These results have important ecological implications. First, the lowest activities of phytoplankton APs at very low pH (5.5) verified that algal AP is sensitive to pH decline and cannot function well at low pH. Although only dinoflagellates were included in the study, this is likely true for other groups of phytoplankton. Second, the differential optimal pH for AP activity among phytoplankton species suggest that their capacities to scavenge DOP for P nutrition will be influenced by OA to different extents. Based on the current trend of CO_2_ increases, pH in future oceans is projected to be about 7.8-7.7 by the end of this century (Orr et al. 2005, Caldeira and Wickett 2005, Doney et al. 2009). Under this scenario, species such as *F. kawagutii* and *P. cordatum* may benefit from OA in terms of utilizing DOP based on AP. In contrast, species such as *A. carterae* and *E. voratum* will have their AP activities undermined by OA. Other species such as *K. mikimotoi* may not be so much impacted by OA in terms of AP-based DOP utilization.

A recent study treating isolated cultures and on-deck incubated field population of *Trichodesmium* indicated that OA combined with P-limitation synergistically depressed nitrogen fixation by this diazotroph (Zhang et al. 2022). Transcriptomic analysis showed that AP expression was upregulated under the OA-P limitation condition, but due to the lack of information on optimal pH for this species’ AP, whether AP activity will be compromised by OA is unclear. More research is needed to dissect how OA impacts P nutrition in phytoplankton and affects physiologies (e.g. N_2_ fixation) and growth in different species.

Finally, our findings suggest that OA combined with P limitation has the potential to alter phytoplankton community structure. Because OA differentially influences phytoplankton species in photosynthetic carbon fixation (depending in part on whether the species has CCM) and DOP utilization, growth rate under OA and P limitation conditions will vary among phytoplankton species. Further research is needed, particularly using mesocosm with mixed species or natural assemblages, to investigate how phytoplankton communities are shaped by OA in combination with P limitation.

## Acknowledgements

The authors wish to thank members of the Marine Ecological Genomics group for kind assistance in various ways. S. Lin was in part supported by the Gordon and Betty Moore Foundation (GBMF) grant #4980.01.

## Author Contributions

SL conceptualized the project and wrote and edited the paper. CG executed the experiments and data analysis and wrote the paper. LL provided technical training and advice. XL provided advice on species selection and the overall workflow.

